# ATPγS unbiases kinesin

**DOI:** 10.1101/2025.05.01.651652

**Authors:** Vishakha Karnawat, Algirdas Toleikis, Nicholas J. Carter, Justin Molloy, Robert A. Cross

## Abstract

Kinesin-1 microtubule motors are ATP-fuelled, twin-headed cargo transporters that step processively along microtubules, with a load-dependent directional bias. Here we show using single molecule optical trapping that 1 mM ATPγS, a slowly-hydrolysed analogue, substantially defeats the biasing mechanism, whereas 1 µM ATPγS supports it. Our data argue that kinesin re-registers itself between steps into an Await-Isomerisation (AI) state that engenders both on-axis steps and off-axis missteps, and that missteps can be rescued. In the AI state, ATP or ATPγS is bound but neck-linker docking and nucleotide hydrolysis are inhibited. Load-dependent exit from the AI state establishes hydrolytic competence via catalytic site closure and coupled neck-linker docking, which guides the tethered head to its next on-axis site. By overpopulating the AI state, ATPγS reveals its pivotal role in the biasing mechanism, whose control logic maximises steered diffusion-to-capture of the leading kinesin-1 head under load, at the expense of futile nucleotide turnover.

---

Kinesins are microtubule-based cargo transporters whose activities are central to the self-organisation 1 of eukaryotic cells and organisms. The archetype of the kinesin superfamily is kinesin-1. Kinesin-1 motors haul cargo in 8 nm steps towards the plus ends of microtubules ^2^, using a walking action in which the twin heads of each molecule interact alternately with the microtubule ^3^. This alternate-heads stepping mechanism allows single kinesin-1 molecules to make headway against loads exceeding 7 pN ^4,5^. Certain features of the mechanism of single molecule stepping under load are firmly established. Whilst waiting for ATP to bind, kinesin-1 adopts a one-head-bound (1HB) waiting state in which one of its twin heads is in an apo state and strongly bound to the microtubule, whilst the other (tethered) head is in an ADP state and is prevented from binding to the microtubule ^6–8^. There is evidence that in this ATP-waiting state, the tethered head is also prevented from binding unpolymerised tubulin ^9^. ATP binding to the microtubule-bound head then frees the tethered head to undergo steered diffusion-to-capture and coupled MT-activated ADP release ^7^. Meanwhile, the trail head hydrolyses ATP and releases Pi, converting it to a weak binding K.ADP intermediate that unbinds from the MT and is recovered ^10^,^11^, re-establishing the 1HB ATP-waiting state. For kinesin under load, we have proposed that directional stepping emerges from a competition between tight-coupled forward steps and loose-coupled backslips ^12^. Recently, fast backsteps were reported that may be related ^13,14^. Whilst the evidence for these features of the kinesin-1 mechanism appears firm, other aspects remain controversial, with recent work revisiting the incidence of substeps ^14^, sidesteps ^15^ and inchworming ^16^.

A potentially helpful way to deconstruct the kinesin-1 stepping mechanism is to change its nucleotide fuel. ATPγS is a slowly-hydrolysed analogue in which one of the hydroxyl groups of the terminal (gamma) phosphate is substituted for a thiol. The hydrolysis reaction produces ADP and thiophosphate. ATPγS turnover drives slow directional gliding of MTs in motility assays ^17^ and slow, disorderly but directional stepping of unloaded kinesin dimers ^18^. We show here using single molecule optical trapping that 1 mM ATPγS largely defeats the guidance mechanism that steers on-axis forward stepping of kinesin-1 under load, whilst 1 µM ATPγS does not. This differential provides new insight into the componentry and action of the guidance mechanism.

## Results

### ATPγS drives slow stepping and competes effectively with ATP

We first confirmed that ATPγS binds tightly, drives slow MT gliding and competes effectively with ATP (**Fig. 1a-c**). The MT gliding rate in 1 mM ATPγS is 12 nm s^-1 17^, whereas in 1 mM ATP it is ∼800 nm s^-1^. Single kinesin molecules in ATPγS step at 13 nm s^-1 18^. Under load in the optical trap, ATPγS-driven single molecule stepping superficially resembles ATP-driven stepping, except that it is much slower (**Fig. 1d-g**). Both 1 mM and 1 µM ATPγS drive kinesin to step with ATP-like processivity to a stall force that approximates that in ATP. For kinesin-1, stall force is defined as the force at which the probabilities of forward steps and backsteps are equal.

**Figure 1:**
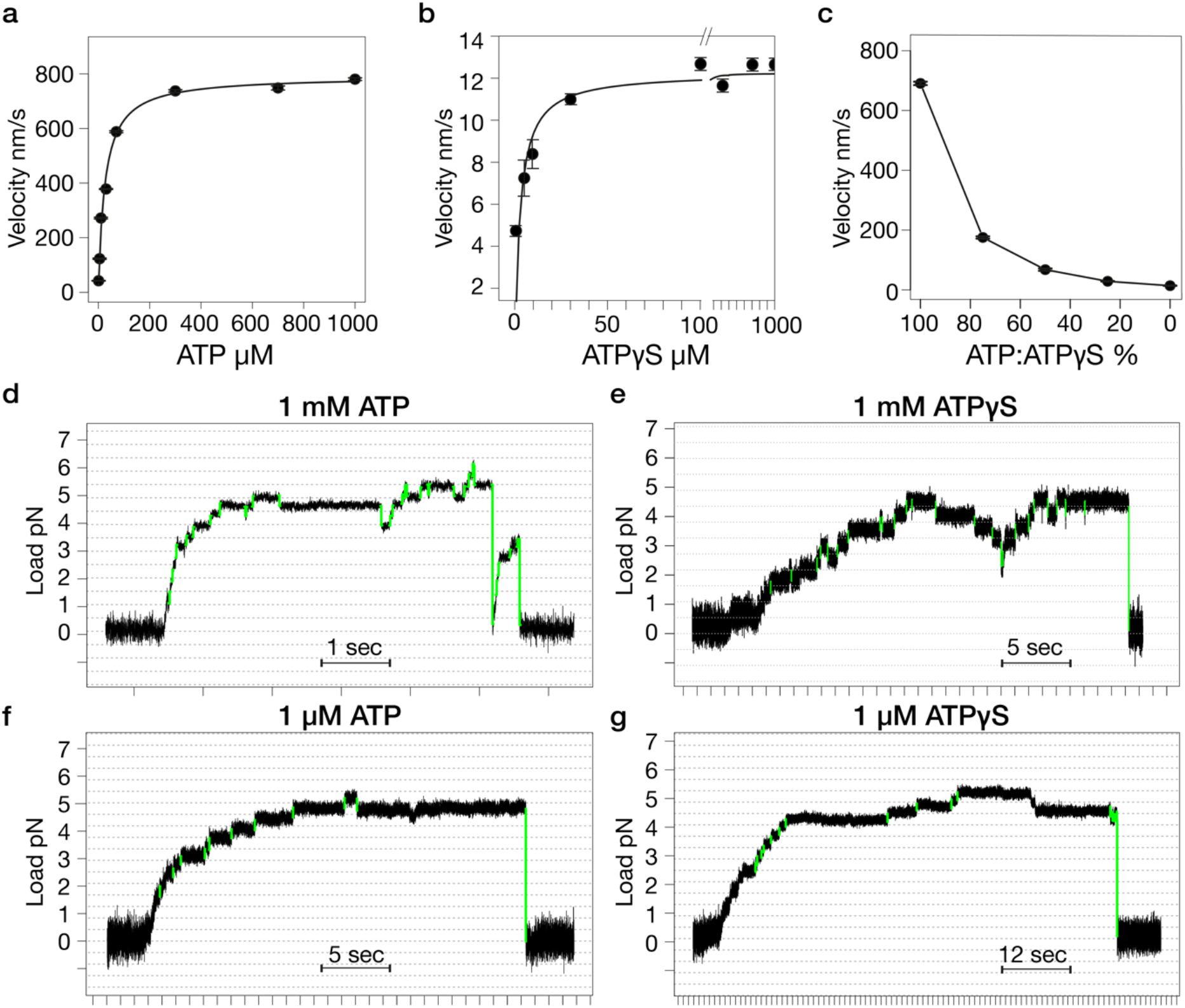
Kinesin motility in ATP and ATPγS. (**a-b**) Microtubule gliding velocity in kinesin surface assays as a function of nucleotide concentration for (**a**) ATP; (**b**) ATPγS. Data are fitted with the Michaelis-Menten function: K_m_ [ATP] = 25.7 μM, V_max_ [ATP] = 0.79 μm/s, K_m_ [ATPγS] = 3.1 μM, and V_max_ [ATPγS] = 0.012 μm/s. (**c**) ERect of the ratio of ATP to ATPγS on microtubule gliding velocity. The x-axis represents the percentage of ATP relative to ATPγS, ranging from 100% ATP (0% ATPγS) to 100% ATPγS, with the corresponding microtubule gliding velocities plotted on the y-axis. Note the ∼75% reduction in gliding velocity with the addition of 25% ATPγS. (**d-g**) example trapping records. Vertical divisions are at 1 second intervals. Horizontal grid lines are 8 nm apart. t-score cutoff (see Methods) for detecting steps in **d** = 15, **e** = 15, **f** = 15, and **g** = 18.

### 1 mM ATPγS substantially defeats the biasing mechanism

The forward to backward (F/B) probability ratio for kinesin stepping gives a measure of the directional bias. In ATP, the F/B ratio diminishes exponentially with load (**Fig. 2a**,**e**). The exponential factors are similar at 1 µM and 1 mM ATP (Compare **Fig. 2e**,**f**), as noted previously for 10 µM and 1 mM ATP ^4,5^. By contrast in 1 mM ATPγS the F/B ratio is only very weakly dependent on load (**Fig. 2c**,**g**). At substall loads in 1 mM ATPγS, the backstep probability within each force bin is increased compared to that in 1 mM ATP, at the expense of forward steps, whilst the probability of longer backslips or detachments is little affected. In both ATP and ATPγS, steps at superstall loads are rare, because detachments are favoured. Using an experimental protocol in which superstall load is rapidly applied by imposing a sudden backwards motion of the microscope stage we were able to improve sampling of kinesin stepping behaviour over a wide range of loads (**Fig. 2i**,**j**). In both 1 mM ATP and 1 mM ATPγS, this did not much change the load-dependence of the F/B step ratio (**Fig. 2k**,**l**).1 mM ATPγS thus substantially, though not entirely, defeats the biasing mechanism that favours forward steps over backsteps under load.

**Figure 2:**
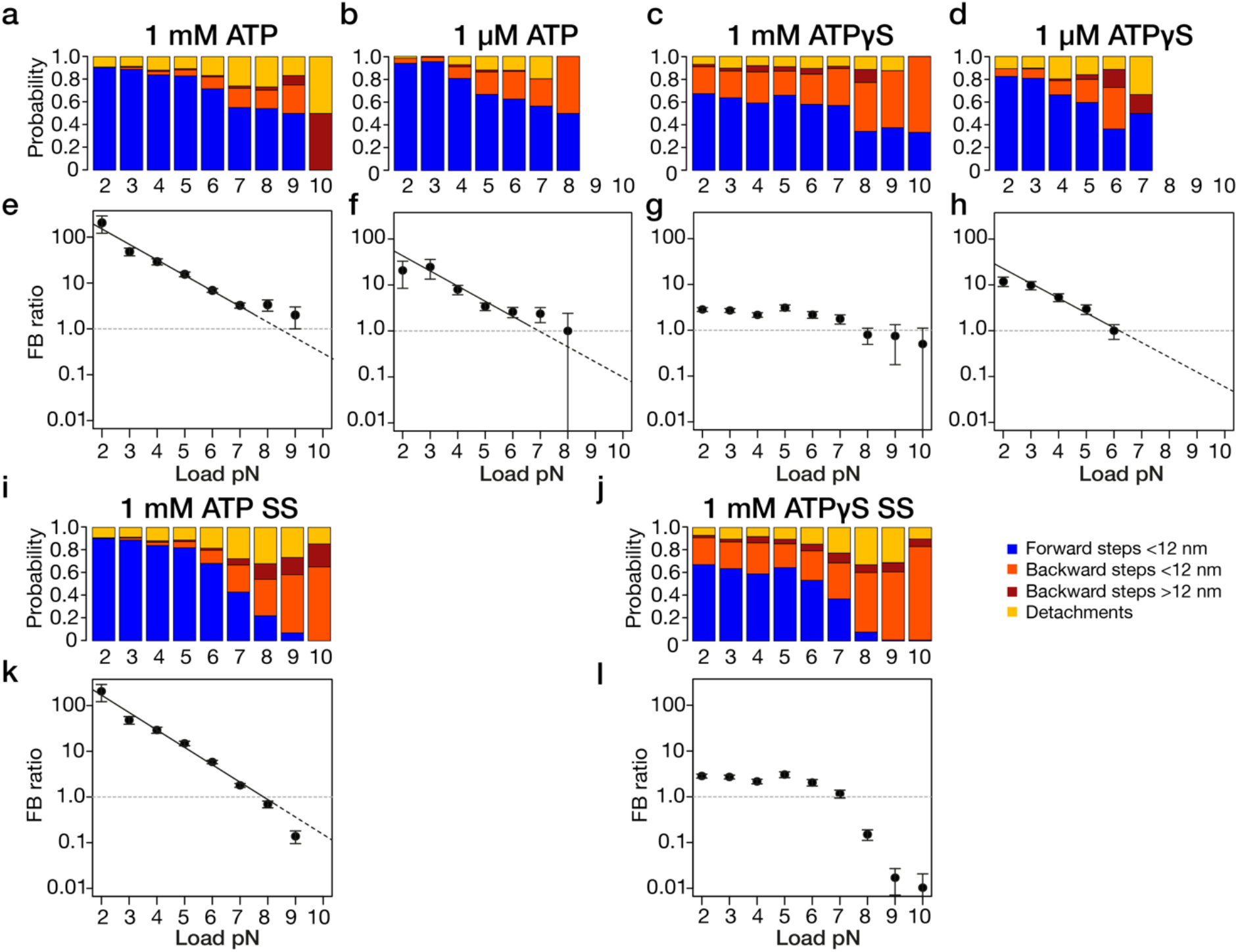
Step-type probabilities and F/B step ratios under load in ATP versus ATPγS. (**a-d**) Probability of forward steps (blue), <12-nm backsteps (orange), >12-nm backsteps (magenta), and detachments (yellow) versus load for kinesin stepping in (**a**) 1 mM ATP, (**b**) 1 μM ATP, (**c**) 1 mM ATPγS, and (**d**) 1 μM ATPγS. (**e-h**) F/B ratio (#forward steps / #backsteps) versus load for the data in **a**-**d**. (**e**) 1 mM ATP (fitted: 2-7 pN), F/B ratio = 713.94 e^-0.78 * load^, (**f**) 1 μM ATP (fitted: 3-6 pN), F/B ratio = 197.23 e^-0.76 * load^, (**g**) 1 mM ATPγS and (**h**) 1 μM ATPγS (fitted: 3-6 pN), F/B ratio = 100.62 e^-0.74 * load^. (**i**,**j**) equivalent to (**a**) and (**b**) except stage was stepped to superstall (SS) force at a trigger load (Methods). (**k**) 1 mM ATP SS (fitted: 2-7 pN), F/B ratio = 977.63 e^-0.88 * load^. (**l)** 1 mM ATPγS SS

### 1 µM ATPγS supports the biasing mechanism

Reducing the concentration of ATPγS to 1 µM largely rescues the exponential load-dependence of the F/B step ratio (**Fig. 2h**). The exponential factor is similar to that in 1 µM ATP and 1 mM ATP (**Fig. 2e**,**f**).

### Forward and backstep dwell times in ATP depend exponentially on load

The dwell time for each kinesin step is the waiting time before each step. To examine the variation of dwell times with load we averaged all dwells within 1 pN force bins at substall loads (**Fig. 3**). For steps in both 1 mM and 1 µM ATP, the mean dwell time within each force bin increases exponentially with load (**Fig. 3a**,**d**), as previously observed, reaching a plateau value at or around stall force.

**Figure 3:**
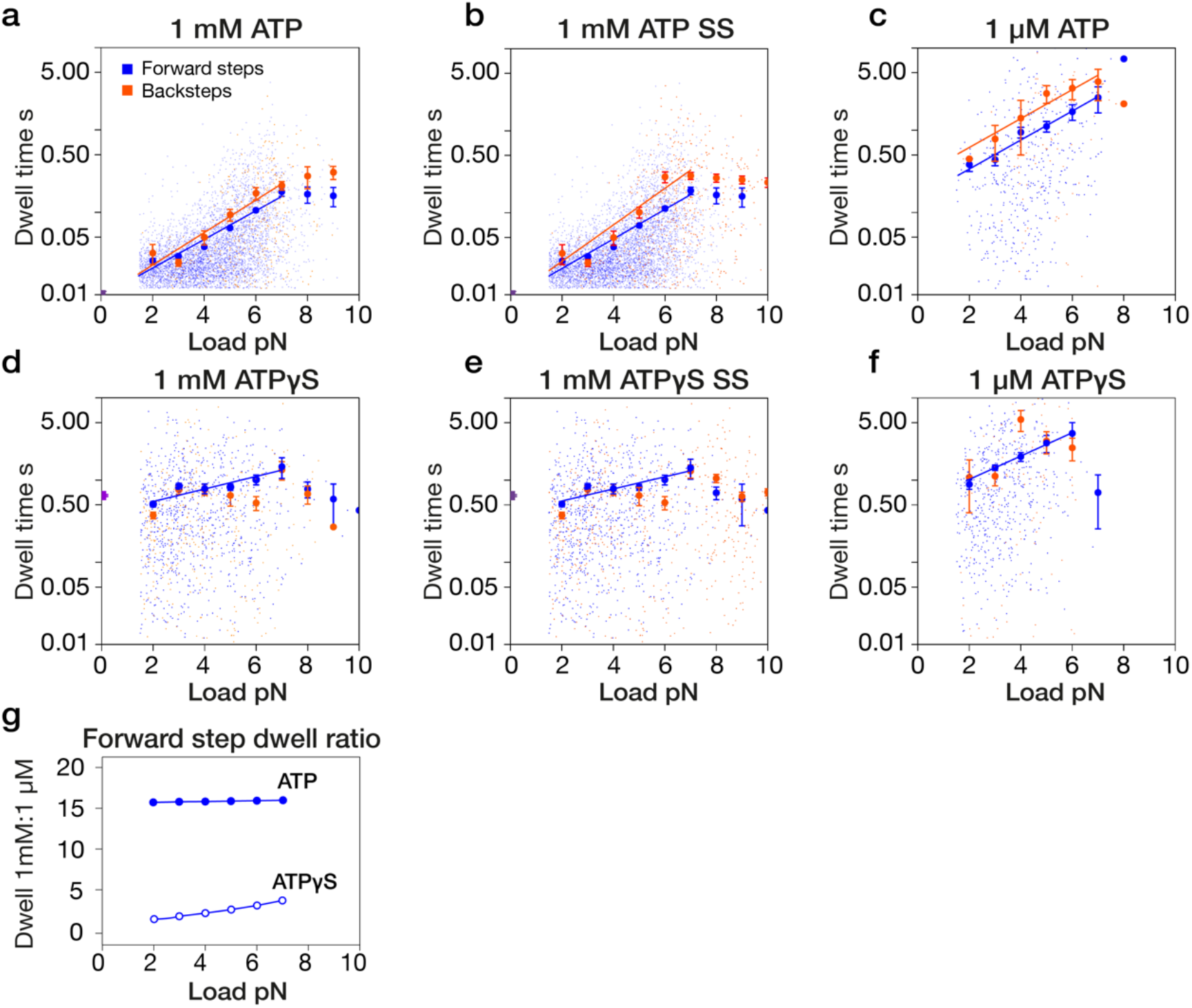
Load-dependence of step dwell times in ATP versus ATPγS. (**a**) Exponential load dependence of step dwell times on load for forward steps (blue) and backward steps (orange) at 1 mM ATP (fitted: 2-7 pN). All the data are plotted. Larger symbols represent the mean dwell times calculated for 1 pN interval bins. Fits are dwell (forward) = 0.0094 e^0.40 * load^ and dwell (backward) = 0.0094 e^0.45 * load^. (**b**) Same but superstall load was applied by triggering a stage-step (Methods). Fits are dwell (forward) = 0.0092 e^0.41 * load^ dwell (backward) = 0.0092 e^0.51 * load^, (fitted: 2-7 pN). (**c**) 1 µM ATP. Fits are dwell (forward) = 0.1500 e^0.41 * load^ and dwell (backward) = 0.2743 e^0.41 * load^ (fitted: 3-7 pN). (**d**) 1 mM ATPγS, dwell (forward) = 0.4067 e^0.17 * load^ (fitted: 2-7 pN). (**e**) Same but applying superstall load by triggering a stage-drag, dwell (forward) = 0.4135 e^0.16 * load^(fitted: 2-7 pN). (**f**) 1 µM ATPγS. Fit is dwell (forward) = 0.5308 e^0.33 * load^ (fitted: 3-6 pN). Fitting in all cases is by least-squares to log (y) = k*x + b. Errors bars are ± standard error (SE). (**g**) Ratios of mean forward step dwell times at 1 µM and 1 mM nucleotide, for ATP and ATPγS.

Further, forward step dwells at any substall load are shorter on average than backward step dwells at the same load (**Fig. 3a**,**d**), consistent with backwards steps originating from a different and later state in the mechanochemical cycle than forwards events ^12^. The ratio of 1 µM vs. 1mM ATP forward step dwell times is invariant with load (**Fig. 3g**).

### Forward and backstep dwell times in 1 mM ATPγS depend only weakly on load

In contrast to our findings with ATP, mean dwell times in 1 mM ATPγS at substall loads are almost load-independent. Whereas dwell times in ATP increase ∼35-fold from around 13 ms at zero load (by extrapolation) to around 400 ms at stall force (**Fig. 3a**), in 1 mM ATPγS, dwells increase only ∼2-fold from ∼500 ms at zero load, to ∼1000 ms at 6 pN (**Fig. 3d**). In 1 mM ATPγS at any particular load, forward and backwards step dwell times are not obviously different from one another.

### Forward and backstep dwell times in 1 µM ATPγS depend exponentially on load

Strikingly, reducing the concentration of ATPγS to 1 µM, well below the apparent K_m_ of 3.1 μM measured in microtubule gliding assays (**Fig. 1b**) restores an exponential load-dependence of dwell times (**Fig. 3f**), with a similar exponential factor to that in 1 µM ATP (**Fig. 3c**). As in 1 mM ATPγS, forward and backstep dwell times at any particular substall load are not obviously different from one another (**Fig. 3d**,**e**). The ratio of forward step dwell times in 1 µM versus 1 mM ATPγS increases with load (**Fig. 3g**).

### A population of aberrantly-long dwells is expanded in ATPγS

To test for any deviation from a single exponential dependence of mean dwell time on load, we made cumulative probability plots ^13^ (**Fig. 4a**). For both forwards and backwards stepping in 1 µM ATP, 1 mM ATP and 1 µM ATPγS, dwell time distributions within each force bin are well fit by single exponentials. Plotting the dwell times predicted by these single exponential fits (**Fig. 4a-d**) confirms the exponential dependence of dwell times on load at substall loads. In both 1 mM and 1 µM ATP, backstep dwell times are consistently longer than forward step dwell times, and the two vary in parallel. In 1 mM ATPγS, dwell times are weakly exponentially dependent on load (**Fig. 4b**). In 1 µM ATPγS, the exponential load-dependence is more marked (**Fig. 4d**), resembling that in 1 µM ATP (**Fig. 4c**). In both 1 µM and 1 mM ATPγS, dwell times for forwards and backwards displacements are not obviously different (**Fig. 4b**,**d**). Within each force bin, a few aberrantly lengthy dwells are inconsistent with the single exponential fit, based on their residuals (Methods). The proportion of these unexpectedly-long dwells increases slightly with load, for both ATP and ATPγS (**Fig. 4e**).

**Figure 4:**
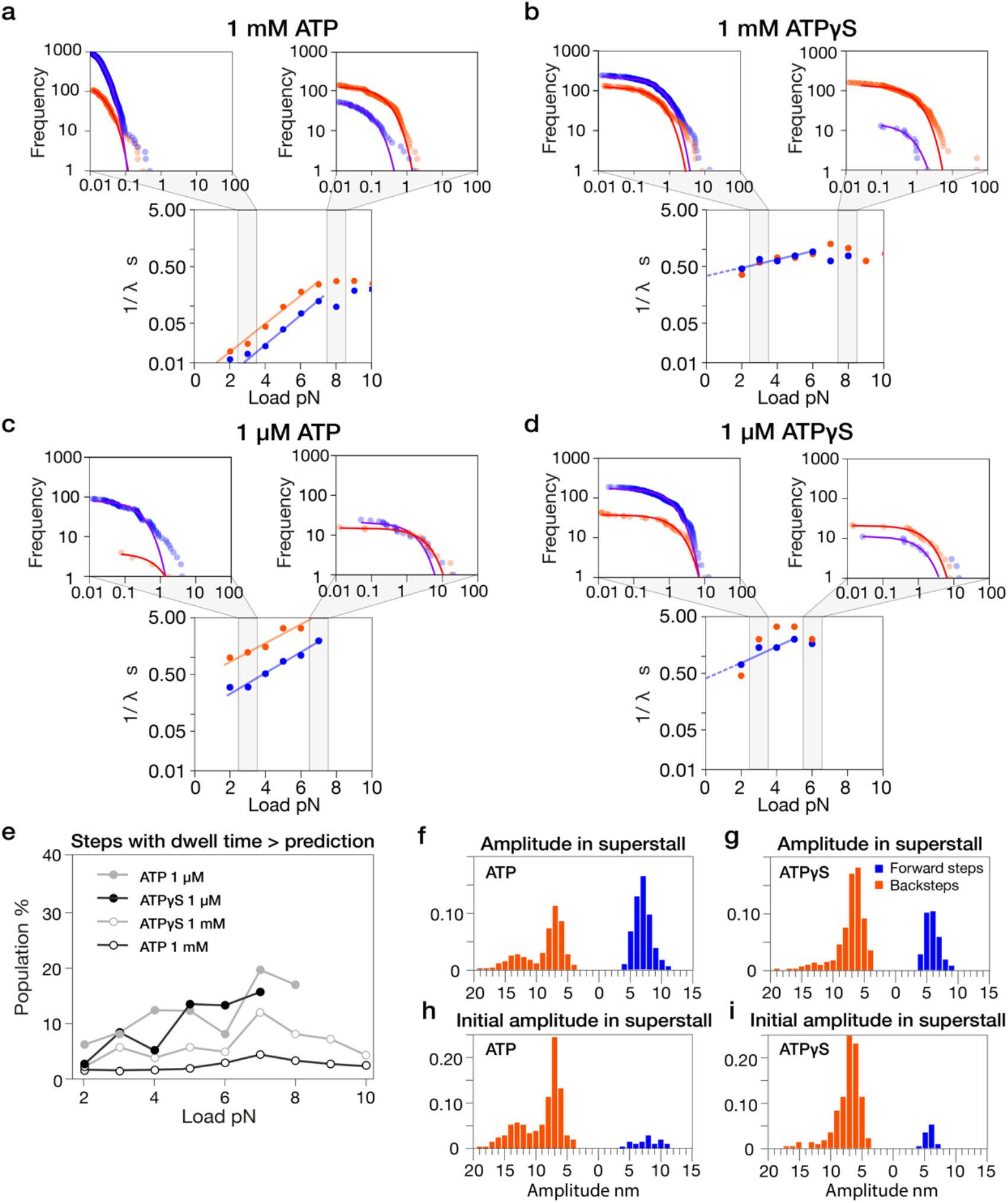
Non-canonical dwells and amplitudes suggest increased misstepping in ATPγS. (**a-d)** Example cumulative density distribution of dwell times of kinesin for forward and backward steps across different conditions, fitted with single exponentials decays, together with load-dependent analysis of fitted dwell times. (**a**) 1 mM ATP, cumulative distribution plots for bins 2.5–3.5 pN (left) and 7.5–8.5 pN (right), with the corresponding plot of mean dwell time (1/λ) vs. load (middle, fitted: 4–7 pN), dwell (forward) = 0.0018 e^0.62 * load^ and dwell (backward) = 0.0049 e^0.58 * load^. (**b**) 1 mM ATPγS, 1/λ vs. load plot (fitted: 2–6 pN) dwell (forward) = 0.3619 e^0.15 * load^. (**c**) 1 μM ATP, 1/λ vs. load plot (fitted: 3-7 pN), dwell (forward) = 0.0971 e^0.43 * load^ and dwell (backward) = 0.3895 e^0.38 * load^. (**d**) 1 µM ATPγS, 1/λ vs. load plot (fitted: 2-5 pN), dwell (forward) = 0.4430 e^0.31 * load^. (**e**) Aberrantly-long dwells as a proportion of the total population of dwells at low and high ATP and ATPγS concentrations. (**f**,**g**) step amplitude distributions for ATP (n = 814) and ATPγS (n = 747) steps obtained at superstall loads after a stage-step. (**h**,**i**) same but only for the initial step that followed each stage-step. n = 213 for **(h)** and n = 233 for **(i)**.

### ATPγS provokes 8 nm backsteps but not larger backsteps

Applying superstall loads of 9 or 10 pN by translating the stage improves sampling at high loads, but often results in a detachment. **Fig. 4f-i** compares amplitudes for these high-load steps, for ATP and ATPγS. We analysed both the set of all high-load steps (**Fig. 4f**,**g**), and the set of first steps obtained following the imposed stage translation (**Fig. 4h**,**i**). In both cases, ATP provokes both 8 nm and 16 nm backwards displacements. ATPγS-driven backwards displacements are almost all 8 nm, pointing to a substantive difference in the mechanics of ATP-versus ATPγS-driven backstepping.

### Supplementing ATPγS with ADP potentiates 8 nm but not larger backsteps

To try to amplify the difference in backstepping behaviour between 1 mM ATPγS and 1 mM ATP, we added ADP. Supplementing 0.9 mM ATP with 0.1 mM ADP increases the incidence of 8 and 16 nm backslips and (especially) full detachments, at the expense of forward steps (**Fig. 5a**,**d**; compare **Fig. 2a**) whilst reducing the exponential dependence of dwell times on load and reducing the difference between the average forward and backstep dwell times at any particular load (**Fig. 5c**,**i**; compare **Fig. 3a, Fig. 4a**),

**Figure 5:**
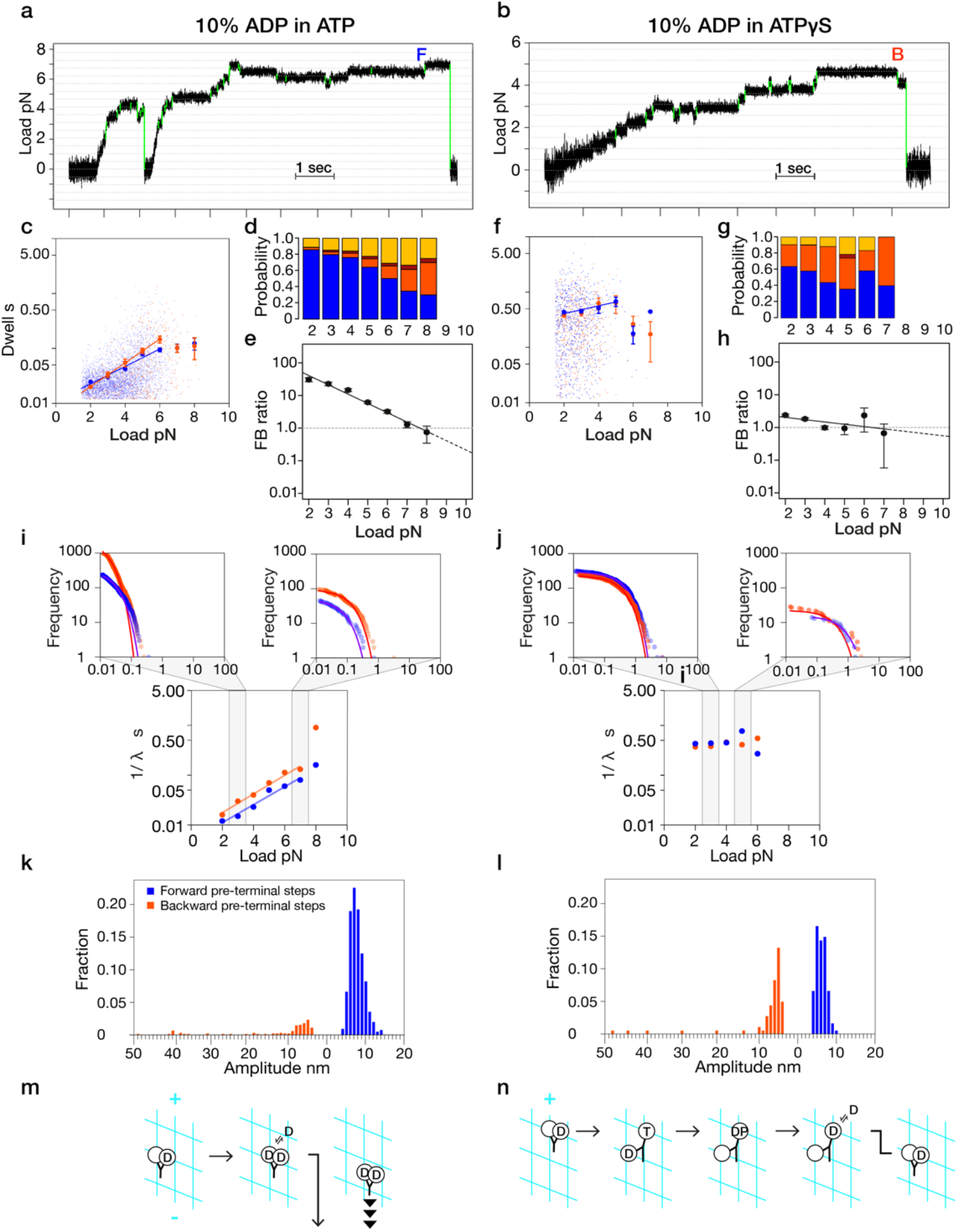
Added ADP causes a pre-terminal 8 nm backstep in ATPγS but not in ATP. Example trapping records for (**a**) 0.9 mM ATP plus 0.1 mM ADP (**b**) 0.9 mM ATPγS plus 0.1 mM ADP. In ATP, detachment is preceded by a forward 8 nm step (F). In ATPγS, detachment is preceded by a backward 8 nm step (B). (**c-h**) Analysis of stepping in 0.9 mM ATP plus 0.1 mM ADP (**c, d, e**) versus 0.9 mM ATPγS plus 0.1 mM ADP (**f, g, h**). (**c, f**) Exponential load dependence of step dwell times. (**c**) Fits are dwell (forward, blue) = 0.0109 e^0.37 * load^ and dwell (backward, orange) = 0.0075 e^0.49 * load^ (fitted: 2-6 pN). (**f**) F<u>its are</u> dwell (forward) = 0.2733 e^0.18 * load^ (fitted: 3-7 pN). (**d, g**) Step-type probabilities (colour coded as in **Fig. 2a–d**). (e, h) Forward-to-backward (F/B) step ratios. (**e**) 10% ADP: ATP (fitted: 2-8 pN), F/B ratio= 152.51 e^-0.66 * load^. (**h**) 10% ADP: ATPγS (fitted: 2-7 pN), F/B ratio = 2.81 e^-0.16 * load^. Fitting in all cases is by least-squares to log (y) = k*x + b. Errors bars are ± standard error (SE). (**i**,**j**) Dwell versus load obtained by fitting cumulative density plots with single exponentials (Methods). Example plots and fits are shown. (**i**) 0.9 mM ATP plus 0.1 mM ADP (left: 2.5–3.5 pN bin; right: 6.5–7.5 pN bin; middle: mean dwell time (1/λ) vs. load), fit: dwell (forward) =0.0049 e^0.41 * load^ and dwell (backward) = 0.0076 e^0.43 * load^. (**j**) 0.9 mM ATPγS plus 0.1 mM ADP. (**k**,**l**) Amplitude distributions for the step immediately preceding detachments in (**k**) ATP (n = 1330), (**l**) ATPγS (n = 181). (**m**,**n**) Potential scenarios for (**m**) detachment without a preterminal tethered backslip. ADP binding directly provokes detachment. (**n**) a preterminal tethered backstep, as in ATPγS. Here, ADP binding increases the probability of a tethered backslip.

By contrast, supplementing 0.9 mM ATPγS with 0.1 mM ADP specifically boosts 8 nm backsteps (**Fig. 5g**; compare **Fig. 2c**), whilst further reducing the already minimal dependence of dwell times on load (**Fig. 5f**,**j**; compare **Fig. 3d, Fig. 4b**). The tendency for added ADP to provoke almost exclusively 8 nm backsteps in ATPγS is directly apparent in trapping records, prior to the detachment events that terminate each processive run (**Fig. 5a**,**b**). In ATP plus ADP, detachments are routinely preceded by a forward step (**Fig. 5k**). By contrast in ATPγS plus ADP, detachments are almost always preceded by an 8 nm backstep (**Fig. 5l**). Terminal backsteps, as a prerequisite for subsequent forward stepping, were previously seen by Guydosh and Block ^6^, in mixtures of ATP and AMPPNP. As we now discuss, terminal 8 nm backsteps in ATPγS may report an obligate transition from a 2HB state to a 1HB state, prior to detachment.

## Discussion

Our data show that 1 mM ATPγS substantially defeats the biasing mechanism for on-axis directional stepping of kinesin-1, whilst making step dwell times longer and almost load-independent. By contrast in 1 µM ATPγS, both directional bias and step dwell time depend exponentially on load, much as they do in ATP. What do these data reveal about the mechanism that biases kinesin stepping under load?

The defining feature of ATPγS, the gamma thiophosphate, is bulkier and potentially more difficult to accommodate within the kinesin active site than the gamma phosphate of ATP ^19^. To account for this and for our findings, we propose that nucleotide binding initially populates an Await-Isomerisation (AI) state (**Fig. 6**) of the kinesin dimer, wherein the trailing head is in a pre-CLOSED conformation in which nucleotide is bound and diffusional excursions of the tethered head are potentiated, but neck-linker (NL) docking on the trailing head is suppressed. In ATP and at low load, we envisage that the pre-CLOSED trail head is in rapid equilibrium with a CLOSED state that is competent for hydrolysis, NL docking, forward stepping (**Fig. 6**, blue pathway) and, following Pi release, for backslipping (**Fig. 6**, orange pathway)^12^. By contrast in ATPγS, and under load in ATP, we propose that isomerisation of the trail head from its pre-CLOSED state to its hydrolysis-competent CLOSED state is retarded, thereby promoting off-axis missteps from the Await-Isomerisation (AI) state (**Fig. 6**, magenta pathway). As we discuss below, this scheme can explain the extra 8 nm backsteps and almost load-independent dwell times seen in 1 mM ATPγS, as well as the apparently more ATP-like properties of 1 µM ATPγS.

**Figure 6:**
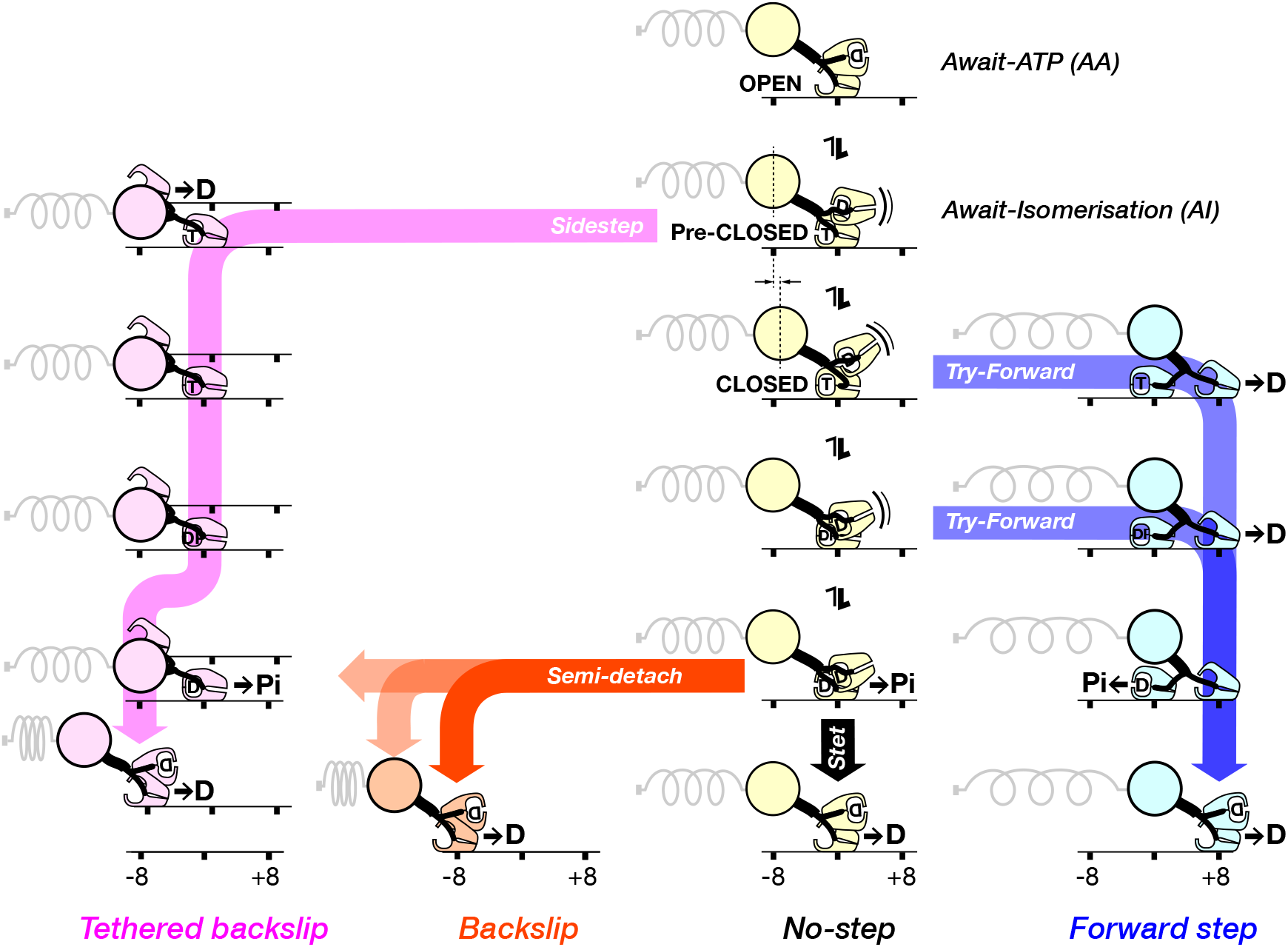
Proposed scheme for kinesin stepping under load. ATP binding to the Await-ATP (AA) state populates the Await-Isomerisation (AI) state, by shifting the trailing, MT-attached head from an OPEN into a pre-CLOSED conformation. This shift triggers the tethered head to begin searching for its next site. In ATP, the AI state is transient at low load, but in ATPγS, the AI state is enriched, because ATPγS retards the isomerisation from pre-CLOSED to CLOSED. Because the AI state is incompetent to dock the NL, it tends to generate missteps, including sidesteps (magenta pathway) that cause the motor to bridge between protofilaments. Bridges can revert directly into the 1HB (one-head-bound) AA state, producing a single 8 nm tethered backslip (as illustrated; favoured in 1 µM ATPγS), or less directly, via multiple rounds of nucleotide turnover that each produce an 8 nm tethered backslip (favoured in 1 mM ATPγS). Following isomerisation of the AI state, the motor enters the Try-Forward era of the cycle, during which only forward steps are possible. NL docking stabilises the CLOSED state, accelerating hydrolysis and steering the tethered head to its next on-axis binding site. Hydrolysis can also proceed via partial NL docking, as shown, at a default slow rate. Competition between this slow default hydrolysis and NL-accelerated hydrolysis creates a load-dependent time window for forward stepping. At high loads, NL docking increasingly fails to occur within this time window, leading to No-steps (yellow pathway) and Backslips (orange pathway). Untethered backslips can evolve into full detachments, but more commonly are rescued via reattachment, reverting the motor to its AA state, allowing forward stepping to be retried, at the expense of futile cycling.

There is firm evidence that NL docking greatly stabilises a hydrolysis-competent CLOSED state of the kinesin head ^20^. Disabling NL docking by mutagenesis profoundly inhibits the MT-activated ATPase of kinesin-1 ^21,22^, indicating that a hydrolytically-competent CLOSED state is stabilised not only by ATP binding and MT binding ^23^, but also by NL docking. Recent cryoEM work ^24,25^ provides direct structural evidence that NL docking can control full closure of the nucleotide pocket, and that AMPPNP can occupy the forward head of a MT-attached dimer, confirming that a kinesin head with an undocked NL can bind nucleotide stably – as in the AI state in our scheme. Very recent data on kinesin--1 ^10^,^11^ further encourage this view. Indeed Niitani et al. argue for a stepping scheme in the absence of external load similar to that which we propose here for the loaded case. In both cases, NL docking is proposed to bias a conformational equilibrium on the trail head that in turn controls both forward stepping and hydrolysis. At high load, NL docking is disfavoured, destabilising the CLOSED state and slowing hydrolysis. Conversely at low load, the NL docks readily, the CLOSED state is stabilised and hydrolysis is promoted. The exponential dependence of dwell time on load (**Fig. 3, Fig. 4b**) then directly reflects the load-sensitivity of the NL docking equilibrium. At very high loads, step dwell times become load-independent, reflecting that at superstall loads, the NL almost never docks, and almost all events are backslips.

In our proposed scheme (**Fig. 6**) ATPγS exerts its effects by stabilising the AI state, thereby promoting stepping without the steering action of NL docking, including sidestepping to a neighbouring PF (magenta pathway), with coupled ADP release. In 1 µM ATPγS, nucleotide binding to the newly-apo head created by ADP release will be unlikely, allowing it to act as a holdfast as its partner progresses through hydrolysis and thiophosphate release into its weak binding K.ADP state (**Fig. 6**, magenta pathway). The partner K.ADP head can then either detach, reverting the motor to its one-head-bound (1HB) AA state (**Fig. 6**, top and bottom rows), or release ADP, or slip back before rebinding and releasing ADP. If it slips back, the length of its slip will be restricted by the combined length of the two extended neck linkers, which act as a tether. Following a tethered backslip, one or more additional ATPγS turnovers by the now-leading head will then allow it to access its ADP state and detach, locking in the tethered backslip and regenerating the1HB AA state (**Fig. 6**, magenta pathway). In 1 µM ATPγS (**Fig. 2h**), the F/B ratio is similar to that in 1 µM ATP (**Fig. 2f**), indicating that forward stepping in 1 µM ATPγS is routinely successful and by implication that off-axis missteps in 1 µM ATPγS are readily rescued. We envisage that following a sidestep and tethered backslip, reversion to the 1HB AA state and hence to the AI state depends on the head that has just released ADP remaining briefly in its apo state, and on the partner head retaining its newly-generated ADP (**Fig. 6**, magenta pathway). These requirements will be met in 1 µM ATPγS, because the probability of ATPγS binding on the timescale of interest will be low. Consistent with this, in 1 µM ATPγS we observe both an exponential load-dependence of dwell times (**Fig. 3f**) and an exponential load-dependence of the F/B ratio (**Fig. 2h**), resembling those seen in 1 µM ATP (**Fig. 2f**). By contrast in 1 mM ATPγS, nucleotide binding is saturated and reversion to the AA and AI states is correspondingly less likely, as indicated by the reduced F/B ratio (**Fig. 2g**) and the reduced load-dependence of dwell times (**Fig. 3d**,**e**). Both effects reflect an excess of backsteps, especially at low load. These backsteps are almost invariably 8 nm (**Fig. 4g**,**i**), consistent with tethered backslips.

Further evidence for tethered backslipping in ATPγS comes from ADP supplementation (**Fig. 5**), which generates extra backsteps that are also almost exclusively 8 nm. We propose that ATPγS causes tethered backslips and that these are preceded by sidesteps from the AI state (**Fig. 5n, Fig. 6**, magenta pathway). Sidestepping by kinesin-1 in 1 mM ATPγS was clearly evident in earlier single molecule tracking of gold nanoparticles attached to kinesin heads ^18^. In ATP at low loads, our scheme predicts that misstepping from the AI state will be rare, because the AI state will exist only very transiently. Accordingly, the motor will tend to track a single protofilament as it partitions between the forward step (blue), no-step (yellow) and backslip (orange) pathways. In ATP we see a spectrum of backwards displacements of 8, 16 or 24 nm, or more (**Fig. 4f**,**h**), indicating that most or all these events are untethered, on-axis backslips ^12^ (**Fig. 6**, orange pathway).

In both ATPγS and ATP, our scheme envisages that nucleotide binds load-independently, but then isomerisation of the AI state feeds the Try-Forward states (**Fig. 6**, blue pathway), for which NL docking is potentiated and load-dependent. Time spent in the AI state thereby becomes indirectly load-dependent and contributes to the exponentially load-dependent dwell times of both forward steps and untethered backslips. In ATP, untethered backslips and detachments (**Fig. 6**, orange pathway) have clearly longer average dwell times than forward steps, because they originate later in the cycle than forward steps ^12^. By contrast in ATPγS, average dwell times for forward and backwards displacements are indistinguishable (**Fig. 3e**). Why is this?

In both 1 µM and 1 mM ATPγS we expect dwell times to be dominated by time spent in the AI state, waiting for the active site to adapt itself to ATPγS. We can estimate this await-isomerisation time to be ∼ 500 ms on average, for both 1 µM and 1 mM ATPγS, by extrapolating load versus dwell time plots to zero load (**Fig. 3d**,**e**,**f**, **Fig. 4b**,**d**). The average step dwell time at any substall load in 1 µM and 1 mM ATPγS will be equal to this await-isomerisation time plus the average dwell time seen in 1 µM or 1 mM ATP, because following exit from the AI state, the rest of the cycle is identical to that in ATP. Indeed, adding ∼500 ms to the dwells for 1 µM and 1 mM ATP does substantially recapitulate the ATPγS data (not shown). A consequence is that the expected slight difference between the average forward and backstep dwell times in ATPγS disappears into the noise.

We believe the scheme in **Fig. 6**, is the simplest that can explain our data. The novel aspect is that each step begins with the motor re-registering itself into its 1HB AI state. This property emerges as a central principle of the biasing mechanism, allowing the motor to pause ATP turnover and wait for tethered diffusion-to-capture, steered by NL docking, to carry the tethered head to its next site. The initial loading of ATP (or ATPγS) to form the AI state directly activates the diffusional search of the tethered head for its next binding site. This feature is indicated by Hackney half-site release ^7^ experiments in solution, in which ATPγS binding to the MT-attached head of a kinesin dimer triggers MT-activated mantADP release from its partner at ∼30 s^-1^, much faster than the ∼2 s^-1^ rate of ATPγS-driven stepping ^18^,^12^,^20^and the ∼3 s^-1^ rate of MT-activated ATPγS turnover (**Fig. 1**). AMPPNP, which is non-hydrolysable on the time scale of interest, also triggers half-site release at ∼30 s^-1 18^, consistent with it creating a pre-hydrolysis (pre-CLOSED) state for which tubulin-binding by the tethered head is potentiated. With ATP loaded and diffusion of the tethered head activated, the equilibrium between the pre-CLOSED and CLOSED states of the trail head shifts dynamically according to the load, causing the average dwell time for forward steps to increase exponentially with the load (**Figs. 3 & 4**). It was previously shown that at zero load, ATPγS-driven kinesin stepping remains processive and plus-end biased, but that steps are more chaotic than in ATP ^18^. It was suggested, because rates of half site release driven by the non-hydrolysable nucleotides AMPPNP and ATPγS are slower than the overall rate of MT-activated ATP turnover, that stepping typically waits for hydrolysis ^26,27^. In our proposed scheme, forward stepping and hydrolysis are coupled, in agreement with these earlier observations, but we envisage that under load, neck linker docking promotes hydrolysis, rather than the other way around. In our scheme, hydrolysis can also occur without NL docking, for example under superstall loads, at a default slow rate that feeds the backslipping (**Fig. 6**, orange) pathway and/or the no-step (**Fig. 6**, yellow) pathway. This rate corresponds to the backslipping rate under superstall load, which is about 2 s^-1^. This slow default rate of ATP hydrolysis might reflect docking of the initial segment of the NL, as shown (**Fig. 6)**. This can substantially accelerate hydrolysis ^21^,^11^.

It is known that when kinesin-1 at zero load collides with an obstacle it usually detaches ^28^, but occasionally steps sideways and continues along a neighbouring PF ^29^. For kinesin-1 under load in ATP, we expect the AI state to be enriched and the probability of off-axis stepping to increase correspondingly. Nonetheless our data imply that in ATP, sidesteps and tethered backslips from the AI state are relatively rare, relative to their incidence in ATPγS. Stepping sideways and backwards at a relatively low ‘leakage’ rate (**Fig. 6**, magenta pathway) might nonetheless be useful in vivo, in situations where a protofilament-level obstacle persistently blocks progress, so that the more usual on-axis backslip-and-retry mechanism (**Fig. 6**, orange pathway) would be ineRective.

We have argued that the mechanics of ATPγS-driven stepping under load reveal an Await-Isomerisation (AI) state of kinesin-1, which engenders not only on-axis forward steps and backslips, but also off-axis missteps. In the AI state, nucleotide is bound and stepping is potentiated but hydrolysis and coupled NL docking are retarded. In the scheme we propose, the AI state is transient at low load in ATP but enriched at high load and in ATPγS. Re-registration of the motor to its AI state at the beginning of each cycle is pivotal to the mechanochemical coupling – it allows NL docking to steer the tethered head to its next on-axis microtubule binding site, with a coupled transition to hydrolytic-competence. When steered diffusion-to-capture of the tethered head fails (for example at high load, or when encountering an obstacle), multiple backslipping pathways lead the motor back via futile turnovers to re-establish its AI state, allowing it to retry forward stepping. The ability to fall back, recover and persistently retry forward stepping under load emerges as a central principle of the biasing mechanism, whose control logic maximises processive forward progress under load by adaptively blending tight-coupled forward steps with loose-coupled no-steps, sidesteps and backslips, at the expense of futile cycling.

## Author Contributions

V.K. performed all the experiments and analyzed all the data. N.J.C. designed and built the optical trap. V.K., A.T., N.J.C., J.R.M and R.A.C. collaborated to design and interpret experiments and write the manuscript.

## Acknowledgements

This work was funded by a Wellcome Investigator Award to R.A.C., grant number 220387/Z/20/Z.

## Data Availability

The data analysed in this study are available from the corresponding author upon request.

## MATERIALS AND METHODS

### Kinesin beads

Unmodified 560-nm polystyrene beads (Polysciences, Warrington, PA) were incubated with purified recombinant full-length *Drosophila* kinesin-1 (12) and diluted stepwise until approximately one-third of the beads displayed motility. Experiments were carried out in BRB80 buffer (80 mM K-PIPES, pH 7.0, 2 mM MgSO_4_, 1 mM EGTA, 3 g/L glucose) supplemented with 1 mM ATP, a glucose-catalase oxygen scavenging system, and 10 mM Taxol.

### Microtubule Preparation

Tubulin from porcine brain was diluted in 1× BRB80 buffer supplemented with 1 mM GTP to a final concentration of 20 µM. The solution was centrifuged at 90,000 rpm for 10 minutes, and the supernatant was incubated at 37°C for 20 minutes to facilitate polymerization. Taxol was then added in a stepwise manner: first, 1.75 µL of 2 mM Taxol was added (final concentration ∼ 75 μM), and the mixture was incubated at 37°C for 10 minutes, followed by the addition of 1 µL of 2 mM Taxol (final concentration ∼ 116 μM) and further incubation at 37°C for 20 minutes. The microtubules were then maintained at room temperature for several hours before use in optical trap assays.

### Flow cells

The flow cell surface was passivated with 0.1 mg/mL casein (SuSoS AG, D€ubendorf, Switzerland). MTs were covalently attached to the coverslip surface using mild glutaraldehyde cross-linking to the 3-Aminopropyl)triethoxysilane (APTES)-silanized surface (21).

### Optical trapping

A custom-built optical trap (15) was used, equipped with a 3-W Nd:YAG 1064-nm laser (IE Optomech, Newnham, England). MTs were initially visualized by differential interference contrast microscopy, and beads were moved into position above the MTs by steering the trap. Imaging was then switched to amplitude contrast, and the image was projected onto the quadrant photodiode detector. Data were recorded at 20 kHz, and the moving average was filtered to 1 kHz during analysis.

Superstall experiments were performed as previously described by Carter and Cross. Briefly, once kinesin reached a 3.5-pN trigger point (2.5 pN when ATPγS was used), the motor was subjected to a predefined high-force load by moving the microtubule (via a piezoelectric stage) toward the plus end. A transient, software-based force-feedback system was employed, utilizing both on-axis and oR-axis bead position data while accounting for trap stiffness, microtubule orientation, and polarity. Microtubule displacement was typically completed within 200 ms, after which the feedback system was deactivated, and the trap and stage positions were held fixed.

### Data analysis

Data were analyzed in R using custom-written code (available on request). Automated step-detection was implemented using t-test analysis. In the t-test analysis, twelve data points before the suspected step and twelve after the step were compared by t-test. Dwell times were defined as the waiting time between two consecutive steps. To ensure the accuracy of dwell time measurements, dwells in each force bin were manually verified. This verification involved visually inspecting the data to identify any missed steps between consecutive dwell times. In cases where a missed step was detected, the corresponding dwell was removed, as it represented a false dwell caused by the missed step. This manual inspection helped eliminate any false dwell times, ensuring the reliability of the data. Only steps above 2 pN could reliably be detected, and only these were processed. For forward step/backstep ratio measurements, the force range 3–8 pN was analyzed to ensure a suRicient number of both forward steps and backsteps. Each force bin includes data at the force shown ± 0.5 pN.

### Dwell Time Analysis and Model Fitting (Figure 4)

The plot represents the dwell time and its frequency for kinesin motor forward or backward steps within the specified force range (pN). Dwell time is displayed on the x-axis using a log scale, while the y-axis shows the frequency of each dwell time occurrence, also on a log scale. The frequency values correspond to the count of occurrences for each dwell time, arranged in descending order. To analyse the dwell time distribution, the data were fitted to a single exponential decay model of the form y = A e ^(−λ * x)^, where A is the amplitude and b is the decay constant. The mean dwell time was calculated as the inverse of the decay constant (1/λ) and reported in seconds for each force bin. The obtained mean dwell time was then plotted against load, and the data were fitted using least-squares regression to the equation log(y) = k*x + b.

### Population with Dwell Times Greater than Predicted Values

To calculate the percentage of dwell times greater than the predicted values, a comparison was made between the observed dwell times and the predicted values generated from the exponential decay model (data in Figure 4). For each dwell time, it was determined whether it exceeded the corresponding predicted value. The number of dwell times greater than the predicted values was counted, and this count was then divided by the total number of dwell times. This result was multiplied by 100 to express the proportion of dwell times that exceeded the predicted values as a percentage.

